# Mapping mouse hippocampal output circuits using direction-selective anterograde transsynaptic transduction

**DOI:** 10.64898/2026.05.13.724828

**Authors:** Taichi Kawamura, Rajeevkumar Raveendran Nair, Ken-Ichiro Tsutsui, Shinya Ohara

**Affiliations:** Laboratory of Systems Neuroscience, Tohoku University Graduate School of Life Sciences, Sendai, Japan; Kavli institute for Systems Neuroscience, NTNU Norwegian University of Science and Technology, Trondheim, Norway; Division for the Establishment of Frontier Sciences, Organization for Advanced Studies, Tohoku University

## Abstract

The hippocampus and its projection targets constitute hippocampal output circuits that are essential for memory, navigation, and emotional regulation. Although cortical and subcortical regions receiving hippocampal projections have been well characterized, it remains unclear how hippocampal signals are processed within these projection target regions. To precisely understand information processing in hippocampal output circuits, it is therefore important to gain genetic access to neuronal subpopulations receiving direct hippocampal inputs for structural and functional analyses. Anterograde transsynaptic transduction using adeno-associated virus (AAV) serotype 1 has emerged as a powerful approach for targeting postsynaptic neurons, but its application is limited by unintended retrograde transport, which leads to false-positive labeling in reciprocally connected circuits. Here, we developed a direction-selective anterograde transsynaptic transduction by combining AAV vectors with distinct infection properties and an intersectional gene expression system. Using the hippocampal-entorhinal circuit, we identified an optimal viral combination that enables predominantly anterograde transsynaptic labeling. We further applied this method to map hippocampal projections to reciprocally connected regions, including the amygdala, providing a robust approach for dissecting complex hippocampal output circuits.

## Introduction

The hippocampus sends information to cortical and subcortical regions through projections from the CA1 region and the subiculum, innervating subsets of neurons within the target regions. These hippocampal output circuits, composed of hippocampal projections and their postsynaptic targets, are thought to underlie a wide range of functions, including memory consolidation, spatial navigation, goal-directed behavior, emotional regulation, and stress responses (Eichenbaum 2017; Fanselow and Dong 2010; Tonegawa, Morrissey, and Kitamura 2018). Previous anatomical studies have investigated the organization of the hippocampal projections using various anterograde tracers, such as phaseolus vulgaris leucoagglutinin (PHA-L), biotinylated dextran amine (BDA), or adeno-associated viral (AAV) vectors (Bienkowski et al. 2018; Cenquizca and Swanson 2007; Jay and Witter 1991; Kloosterman, Witter, and Van Haeften 2003). Although these studies have revealed the diverse projection patterns of the hippocampus, it remains unclear how hippocampal information is processed within targeted regions at the level of defined postsynaptic cell populations. To precisely understand information processing in hippocampal output circuits, it is therefore essential to identify the neural subpopulations within the target regions that directly receive hippocampal inputs.

One powerful approach to identify such target subpopulations is anterograde transsynaptic tracing, which enables the labeling of postsynaptic neurons receiving direct projections from neurons at the injection site. Recently, AAV serotype 1 (AAV1) has emerged as a powerful tool for this purpose, as it can transsynaptically spread to postsynaptic neurons through synaptic vesicle release when injected at high titer (Zingg et al. 2017, 2020). Using this approach, we recently identified a population of neurons in medial frontal cortex (MFC), commonly referred to as the medial prefrontal cortex, that receive direct hippocampal inputs (Alemán-Andrade et al. 2025). This method enabled a comprehensive analysis of the hippocampal-MFC circuit, including the laminar distribution and cell-type identification of target MFC neurons.

However, unlike the hippocampal–MFC circuit, many cortical regions are reciprocally connected with the hippocampus. A major limitation of AAV1-based transsynaptic tracing is that it cannot be applied to such reciprocally connected circuits. This is because high-titer AAV1 also undergoes retrograde transport, resulting in the labeling not only of postsynaptic neurons via anterograde transsynaptic spread, but also of neurons projecting back to the injection site. Consequently, this approach can produce false-positive labeling and cannot accurately identify hippocampal target neurons in reciprocally connected regions such as the entorhinal cortex (EC) and amygdala (AMG).

In this study, we sought to improve this viral strategy to investigate how hippocampal outputs target reciprocally connected regions. To this end, we combined AAV vectors with distinct infection properties with the INTRSECT system, an intersectional gene expression system that utilizes multiple recombinases (Fenno et al. 2014, 2020), and developed an intersectional viral strategy, termed direction-selective anterograde transsynaptic transduction (dsATT). We evaluated this method in a well-characterized hippocampal-entorhinal circuit (Ohara et al. 2023; Witter et al. 2017), and identified an optimal AAV combination that enables predominantly anterograde transsynaptic labeling. We then applied this method to dissect various hippocampal targets including the amygdala which forms reciprocal connections with the ventral subiculum. This approach provides a powerful tool to investigate the neural basis of complex brain functions that arise from reciprocally connected networks.

## Methods

### Animals

Adult male and female C57BL/6N mice, which were purchased from Japan SLC (Shizuoka, Japan), were used in this study. Animals were group housed at a 12:12 hr reversed day/night cycle and had ad libitum access to food and water. All experiments were approved by the Center for Laboratory Animal Research, Tohoku University (Projects: 2019LsA-007). The experiments were conducted in accordance with the Tohoku University Guidelines for Animal Care and Use.

### Surgical procedures and virus injections

Animals were anesthetized with isoflurane in an induction chamber and were injected intraperitoneally with ketamine (80 mg/kg) and xylazine (10 mg/kg). Mice were mounted in a stereotaxic frame, and deep anesthesia was maintained throughout the operation by mask inhalation of isoflurane at concentrations between 1.5–2.5%. Eye ointment was applied to the eyes of the animal to protect the corneas from drying out. The skin overlying the skull was disinfected with iodide, and local anesthesia (lidocaine, 10 mg/kg) was injected subcutaneously. The skull was exposed with a rostrocaudal incision, and a small burr hole was drilled above the injection site. The coordinates of the injection sites are as follows (anterior to either bregma (APb) or transverse sinus (APt), lateral to sagittal sinus (ML), ventral to dura (DV) in mm): dorsal hippocampal CA1 region (dCA1;APb - 1.9, ML 1.3, DV 1.4), ventral hippocampal CA1 region (vCA1; APt +1.9, ML 3.6, DV 2.5), ventral subiculum (vSUB; APt +1.6, ML 3.6, DV 2.5), dorsal medial entorhinal cortex (MEC; APt +0.5, ML 3.4, DV 1.9), amygdala (AMG; APb -1.6, ML 3.0, DV 4.8). Viruses were injected into the target areas by pressure injection at the rate of 10 nl per minute using a glass micropipette (tip diameter = 20–30 µm) connected to a 1 µl Hamilton microliter syringe. Following each injection, the pipette was left in place for another 10 minutes before being withdrawn. The wound was sutured and the animal was monitored for recovery from anesthesia, after which it was returned to its home cage.

To test the Con/Foff construct, which shows Cre-recombinase-dependent expression that is suppressed by Flpo recombinase, 100–200 nl of AAV cocktail, constituting of AAV1-hSyn-Cre-WPRE (1.05 × 10^13^ GC/mL, 75–100 nl, Addgene #105553) and AAVrg-EF1α- mCherry-IRES-Flpo (1.1 × 10^13^ GC/mL, 75–100 nl, Addgene #55634), was injected into the dorsal CA1. In the same animal, AAV with Con/Foff construct (AAV8-EF1α- Con/Foff2.0-EYFP, 2.1 × 10^13^ GC/mL, 100 nl, Addgene #137162) was injected into the dorsal MEC (n = 5).

To test the Coff/Fon construct, which shows Flpo-recombinase-dependent expression that is suppressed by Cre recombinase, 100 nl of AAV cocktail, constituting of AAV-EF1α-Flpo (1.4 × 10^13^ GC/mL, 50 nl, Addgene #55637) and AAVrg-hSyn-Cre-WPRE (3.9 × 10^12^ GC/mL, 50 nl, Addgene #105553), was injected into dorsal CA1, while AAV with Coff/Fon construct (AAVDJ-hSyn-Coff/Fon EYFP-WPRE, 3.4 × 10^12^ GC/mL, 100 nl, UNC vector core) was injected into the dorsal MEC (n = 5). The same viral combination was used to examine the organization of the vCA1-MEC circuit (n = 4) and the vSUB-AMG circuit (n =5).

To analyze the hippocampal output circuit receiving inputs from vSUB at whole-brain level, we systemically delivered a VTKS6 construct (Pouchelon et al. 2022) using the blood-brain barrier-penetrating capsid, AAV-PHP.eB (Chan et al. 2017) (n = 3). The VTKS6 construct enables Flpo recombinase–dependent expression that is suppressed by Cre recombinase. Mice that had received recombinase-expressing AAVs in vSUB were deeply anesthetized with isoflurane, and ophthalmic anesthetic (4% lidocaine) was applied to the right eye. AAV-PHP.eB-VTKS6-GFP (6.47 × 10^13^ Vg/ml, Addgene #178714; 25 μl) was mixed with 100 μl of saline, and the entire volume (125 μl) was injected into the venous sinus of the right eye using a 30-gauge insulin needle. VTKS6 was selected over Coff/Fon construct because it enabled robust amplification of the GFP signal using commercially available polyclonal antibodies (see below for details).

As an anatomical reference for vSUB projection patterns, anterograde and retrograde viral tracing experiments were also performed (n = 4). 50–60 nl cocktail of anterograde and retrograde AAVs, constituting of AAV1-CaMKIIα-mCherry (2.8 × 10^12^ GC/mL, Addgene #114469) or AAV9-hSyn-mCherry (1.0 × 10^13^ GC/mL, Addgene #114472) and AAVrg-hSyn-EGFP, (0.7–1.6 × 10^13^ GC/mL, Addgene #50465), was injected into vSUB. The titer of AAV1 was selected to minimize unintended anterograde transsynaptic or retrograde labeling, as the efficiency of transsynaptic spread strongly depends on viral titer (Zingg et al. 2017). Under this experimental condition, no labeling indicative of unintended transsynaptic or retrograde spread was observed at a level that would affect subsequent histological analysis.

### Immunohistochemistry and imaging

After two to three weeks of survival, the injected mice were euthanized with isoflurane, and subsequently transcardially perfused, first with Ringer’s solution (0.85% NaCl, 0.025% KCl, 0.02% NaHCO_3_) and then with 4% paraformaldehyde (PFA) in 0.1 M phosphate buffer (PB). Brains were removed from the skull, post-fixed in PFA for four hours, and put in a cryo-protective solution containing 20% glycerol and 2% dimethylsulfoxide (DMSO) diluted in 0.125 M PB. A freezing microtome was used to cut the brains into 45-µm-thick sections in either the coronal or sagittal plane, which were collected in six equally spaced series for processing.

To visualize Cre, sections were stained with primary antibodies (1:2000, mouse anti Cre, Millipore #MAB3120; 1:2000, rabbit anti-Cre, Novagen #69050) and secondary antibodies (1:400, Alexa Fluor 647 goat anti-mouse IgG, Jackson ImmunoResearch #115-605-146; 1:400, Cy3 goat anti-rabbit IgG, Jackson ImmunoResearch #111-165-144), while Flpo, which was co-expressed with mCherry, was visualized with a primary (1:500, rabbit anti-DsRed, Clontech #632496) and secondary antibodies (1:400, Cy3 goat anti-rabbit IgG, Jackson ImmunoResearch #111-165-144).

GFP signal was enhanced with primary antibodies (1:1000, chicken anti-GFP, abcam ab13970; 1:1500, rabbit anti-GFP, Invitrogen #A11122) and secondary antibodies (1:400, Alexa Fluor 488 goat anti-chicken IgG, Thermo Fisher Scientific #A11039; 1:400, Alexa Fluor 647 goat anti-chicken IgG, Thermo Fisher Scientific #A21449; 1:400, Cy3 goat anti-rabbit IgG). The same antibodies were used to amplify VTKS6-GFP signals. On the other hand, a custom monoclonal antibody recognizing EYFP amino acids 132–149 (Cosmo Bio Co., Ltd., Japan) was used to selectively amplify EYFP signals expressed from the Coff/Fon construct.

For delineation purposes, sections were stained with primary (1:1000, guinea pig anti-NeuN, Millipore #ABN90P; 1:1000, mouse anti-NeuN, Millipore #MAB377) and secondary antibodies (DyLight 405 goat anti-guinea pig IgG, Jackson ImmunoResearch #106-475-003; Alexa Fluor 647 goat anti-mouse IgG, Jackson ImmunoResearch #115-605-003).

To identify GABAergic neurons, the sections were stained with a primary (1:200, Anti-GAD67 Antibody, Millipore, #MAB5406) and secondary antibody (1:400, Cy3 goat anti-mouse IgG, Jackson ImmunoResearch # 115-165-003).

For immunofluorescence staining, floating sections were rinsed in phosphate buffered saline (PBS) containing 0.1% Triton X-100 (PBT), followed by a 60 min incubation in blocking solution containing 5% goat serum in PBT at room temperature (RT). Sections were subsequently incubated with primary antibodies diluted in the blocking solution for 20 hr at 4 °C, washed in PBT (3 × 10 min), and incubated with secondary antibodies diluted in PBT for 4–6 hr at RT. For GAD67 immunostaining, all processes were performed without the use of Triton X-100. Finally, sections were washed in PBS (3 × 10 min), mounted on gelatin-coated slides, and coverslipped with Entellan new (Merck Chemicals, #107961).

Sections were imaged using an automated scanner (Zeiss Axio Scan Z1). To quantify the colocalization of GFP-, Cre-, and Flpo (mCherry)-immunolabeling, confocal images were acquired in sections taken at every 270 μm throughout the entorhinal cortex, using a confocal microscope (Zeiss LSM 900 AxioImager Z2) with a 20× objective (Plan Apochromat 20×, NA 0.8, Carl Zeiss). The number of immunohistochemically labelled neurons was quantified in a fixed Z-level of the confocal images.

### Whole-brain quantification of labeling and regional correlation analysis

To examine the relationship between anterograde axonal labeling and anterograde transsynaptic labeling across the brain, regional labeling values were quantified using atlas-based registration. Coronal section images were registered to a common anatomical reference using Atlas-Based Brain Alignment (ABBA) implemented in QuPath (Bankhead et al. 2017). All sections were aligned to the Kim laboratory mouse brain atlas, which provides unified anatomical labeling across the mouse brain (Chon et al. 2019).

For each atlas-defined brain region, two measurements were obtained. First, the intensity of anterogradely labeled axons (ATG label intensity) from vSUB was quantified using anterograde tracing datasets (n = 4). For each section, the median fluorescence intensity across the entire section was used as a background estimate and subtracted from all pixel values to reduce background signal. The mean fluorescence intensity within each atlas-defined region was then calculated and used as a measure of ATG label intensity. Second, the density of anterograde transsynaptic labels (ATG-TS labels) from vSUB was obtained from the whole-brain mapping datasets using anterograde transsynaptic tracing with systemic AAV-PHP.eB delivery (n = 3). Anterograde transsynaptically labeled neurons were detected using the cell detection function in QuPath, and the number of detected labels within each atlas-defined region was divied by the total region area to obtain the density.

For each atlas-defined brain region, the ATG label intensity and the number of transsynaptically labeled neurons were averaged across animals. To ensure reliable regional measurements, regions were included in the analysis only when the average number of labeled neurons exceeded 10 and the mean regional area exceeded 1 mm². Pearson correlation and linear regression analyses were performed to examine the relationship between the ATG label intensity and the density of ATG-TS labels across brain regions. Regions included in the analysis and anatomical abbreviations are listed in Supplementary Table 1.

### Visualization of labeling distribution

To visualize the spatial distribution of labeling around amygdala, section images including amygdala were registered to a common anatomical reference using ABBA implemented with QuPath software (Bankhead et al. 2017). All sections were aligned to the Kim laboratory mouse brain atlas, which provides enhanced and unified anatomical labeling across the mouse brain (Chon et al. 2019).

Atlas-aligned coordinates were aggregated into a three-dimensional voxel grid. The grid resolution was set to 50 μm × 50 μm in the XY plane and 300 μm along the Z axis, reflecting the inter-section sampling interval. Within each grid, either the number of labeled neurons or mean fluorescence intensity associated with labeled axons was calculated. The resulting three-dimensional voxel histograms were smoothed using three-dimensional Gaussian filtering to generate continuous spatial density maps. These density maps were computed in three dimensions and subsequently visualized as two-dimensional coronal cross-sections aligned to the reference atlas. Contour lines indicate upper percentiles of voxel-wise density values and are intended for qualitative visualization of spatial distribution patterns.

Atlas-based analyses were restricted to the ipsilateral side of the injection and limited to a subset of atlas-defined regions surrounding the amygdala. Anatomical nomenclature follows the Kim laboratory mouse brain atlas (Chon et al. 2019), with minor modifications.

## Results

### Conventional anterograde transsynaptic tracing in hippocampal-entorhinal circuit

We first tested conventional anterograde transsynaptic tracing using AAV1 in the hippocampal-entorhinal circuit, which reciprocally connects the hippocampal CA1 with the medial entorhinal cortex (MEC). This connectivity is layer specific: MEC layer III (LIII) neurons project to the CA1, whereas deep-layer neurons, particularly layer Vb (LVb), receive inputs from the CA1 (Ohara et al. 2023; Sürmeli et al. 2015; Witter et al. 2017). Owing to this clear input-output segregation, this circuit provides a useful model to evaluate the directional specificity of anterograde transsynaptic tracing methods.

We injected AAV1 expressing Cre recombinase (AAV1-hSyn-Cre-WPRE) into dorsal CA1 (dCA1) and a Cre-dependent AAV expressing EGFP (AAV8-hSyn-DIO-EGFP) into dorsal MEC (Fig. 1a,b). This led to Cre and EGFP expression in both MEC LVb and LIII (Fig. 1c,d). The labeling observed in LVb likely reflected anterograde transsynaptic transport, whereas the labeling in LIII was likely mediated by retrograde transport. Notably, the intensity of Cre/EGFP-labeling was higher in LIII than in LVb neurons, suggesting that retrograde transport of AAV1 may be more efficient than its anterograde transsynaptic spread. These results demonstrate that conventional AAV1-based anterograde transsynaptic tracing produces substantial off-target labeling due to retrograde transport.

**Fig. 1.**
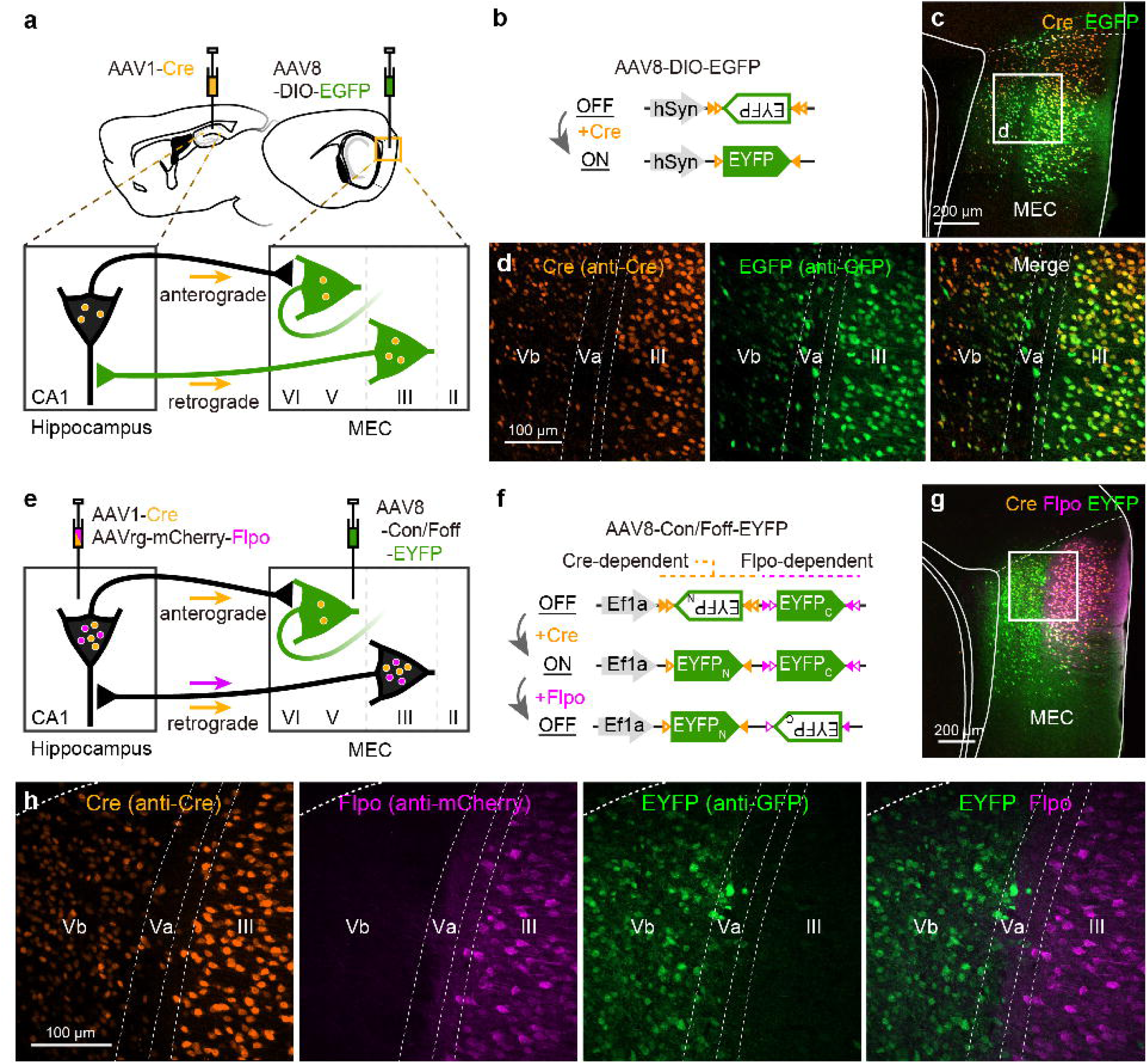
Strategy for achieving anterograde transsynaptic tracing with high direction specificity. **a,b** Schematic of conventional anterograde transsynaptic tracing with AAV1. AAV1-Cre and AAV8-DIO-EGFP were injected into the dorsal hippocampal CA1 and dorsal MEC, respectively. Note that CA1 and MEC are reciprocally connected: MEC layer III neurons project to CA1, whereas layer V/VI neurons receive input from CA1. **c, d** Representative fluorescent micrographs of MEC showing the distribution of neurons expressing Cre (orange) and GFP (green). Cre- and GFP-labeled neurons were observed in both layers Vb and III of MEC, reflecting anterograde transsynaptic labeling and retrograde transport of AAV1, respectively. **e, f** Schematic diagram of the intersectional strategy for direction-selective anterograde transsynaptic transduction (dsATT) using AAV1-Cre, AAVrg-mCherry-Flpo, and AAV8-Con/Foff-EYFP (F). **g, h** Representative fluorescent micrographs of MEC. EYFP labeling driven by the intersectional Con/Foff virus was selectively localized to the deep layers of the dorsal MEC which express Cre but not Flpo.

### Direction-selective anterograde transsynaptic tracing in hippocampal-entorhinal circuit

To overcome this limitation and enable the study of reciprocal circuits, we devised a strategy that combines three different AAVs: AAV1, a retrograde AAV (AAVrg), and an intersectional AAV which expresses EYFP under the presence of specific combination of recombinases (Fenno et al. 2014, 2020).

We co-injected AAVrg expressing Flpo recombinase (AAVrg-EF1α-mCherry-IRES-Flpo) together with AAV1-hSyn-Cre-WPRE into the dCA1 (Fig. 1e). Simultaneously, we injected an intersectional AAV expressing EYFP in the presence of Cre and in the absence of Flpo recombinase (AAV8-EF1α-Con/Foff2.0-EYFP, Fig. 1f) into the dorsal MEC. As in the previous experiment, Cre-labeled neurons were observed in both LVb and LIII of dorsal MEC (Fig. 1g,h). In contrast, Flpo expression —visualized by mCherry—was restricted to superficial MEC layers, consistent with the retrograde specificity of AAVrg. Consequently, EYFP expression was largely confined to LVb neurons, which express Cre but not Flpo, whereas LIII neurons, which express both Cre and Flpo, showed sparse EYFP labeling. This result indicate that the intersectional gene expression strategy is useful for selectively targeting postsynaptic target neurons of the hippocampus while suppressing off-target labeling caused by retrograde infection. Hereafter, we termed this strategy as direction-selective anterograde transsynaptic transduction (dsATT).

### Improve labeling specificity of dsATT

Although the dsATT based on Con/Foff construct enabled selective labeling of MEC deep-layer neurons—the primary recipients of CA1 output—this specificity was limited to the most dorsal portion of MEC. In more ventral MEC, EYFP-labeled neurons were observed in both deep layers and LIII (Fig. 2a,b). Notably, 36.8 ± 5.6 % of EYFP- labeled LIII neurons co-expressed Flpo and mCherry, indicating off-target EYFP expression of the Con/Foff reporter (Fig. 2c). Previous work has shown that Flpo mediates recombination less efficiently than Cre, and that high Flpo expression is required for Con/Foff experiments (Fenno et al. 2020). Because both the number and fluorescent intensity of mCherry-labeled neurons were low in the ventral MEC (Fig. 2b), we hypothesized that insufficient Flpo expression underlies this off-target labeling. To ensure sufficient Flpo expression, we extended the Flpo expression period by injecting the cocktail of recombinase-expressing AAVs into dCA1 15 days prior to the injection of Con/Foff-EYFP into MEC. However, this approach did not improve labeling specificity, as 34 % of EYFP-labeled LIII neurons still showed off-target expression (Supplementary Fig. 1).

**Fig. 2.**
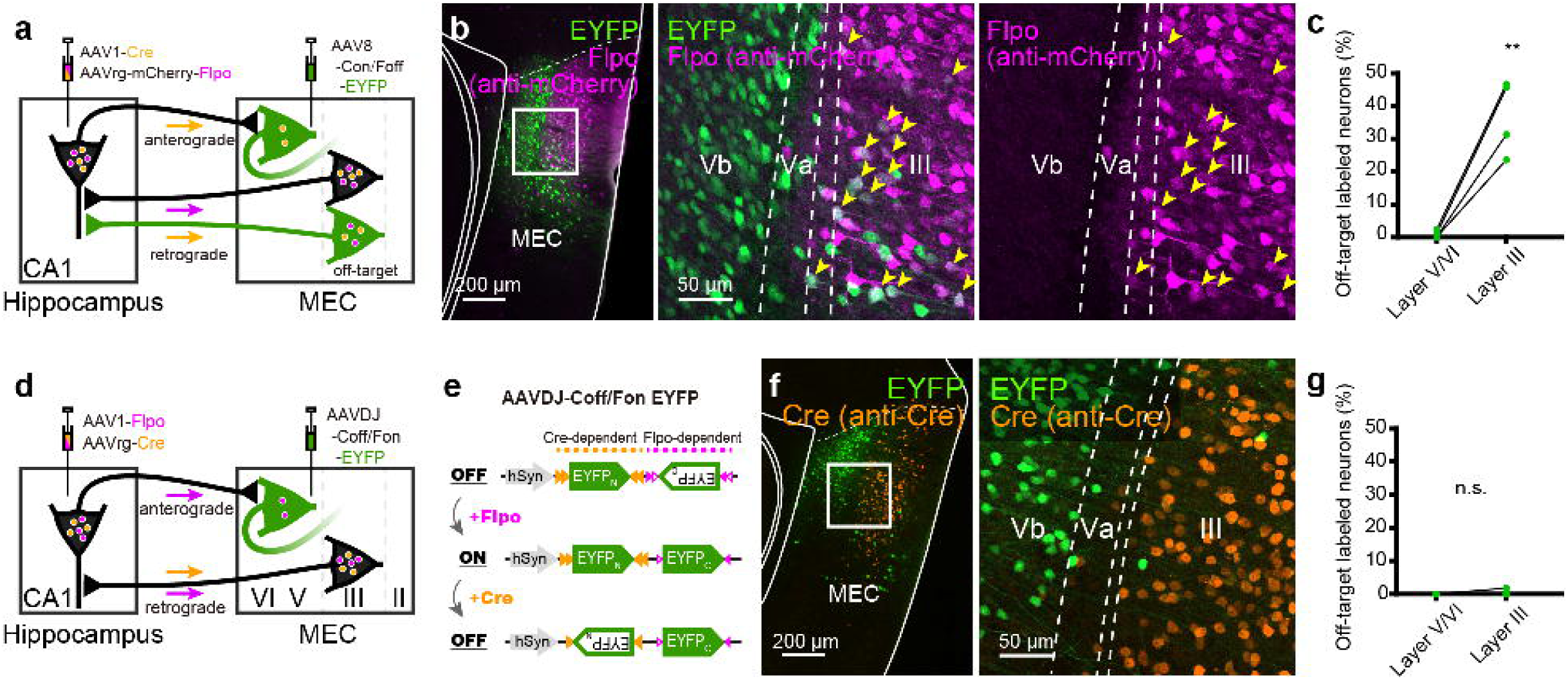
Reduction of off-target labeling using the Coff/Fon construct. **a** Schematic diagram of dsATT using the Con/Foff construct. **b** The representative fluorescent images taken from a sample receiving injections of all three AAVs (AAV1-Cre, AAVrg-mCherry-Flpo, and AAV8-Con/Foff-EYFP), as in Fig. 1b. The white box in the low-magnification image (left) indicates the region shown at higher magnification (right). Yellow arrowheads in the high-magnification image indicate off-target labeled neurons that express both EYFP and mCherry-Flpo. **c** Proportion of off-target labeled neurons in layer III and in deep layers (V/VI) of MEC (N = 4; two-tailed paired t test, t3 = 6.95, ** *p* = 0.006). **d,e** Schematic diagram of dsATT using AAV1-Flpo, AAVrg-Cre, and AAVDJ-Coff/Fon-EYFP. **f** Representative fluorescent image showing the distribution of EYFP- and Cre-expressing neurons in MEC. **g** Proportion of off-target labeled neurons expressing both EYFP and Cre in layer III and in deep layers (V/VI) of MEC (N = 4; two-tailed paired t test, t3 = 1.0, p = 0.39).

We next adopted an alternative strategy using a different intersectional AAV, AAVDJ-hSyn-Coff/Fon EYFP-WPRE, which expresses EYFP in the presence of Flpo but not Cre (Fig. 2d,e; Fenno et al., 2020). Since Cre is a more efficient recombinase than Flpo, this design was expected to reduce false-positive EYFP expression. A high-titer AAV1 expressing Flpo (AAV1-EF1α-Flpo) was injected into the dCA1 together with AAVrg expressing Cre (AAVrg-hSyn-Cre-WPRE), and Coff/Fon-EYFP was simultaneously injected into the dorsal MEC (Fig. 2d). This viral combination markedly improved labeling specificity, and off-target labeling was rarely observed in MEC LIII (Fig. 2f,g). These results demonstrate that Coff/Fon based dsATT substantially enhances the specificity of AAV1-mediated anterograde transsynaptic labeling.

### Analysis of the hippocampal-entorhinal circuits using dsATT

We further assessed the utility of the Coff/Fon-based dsATT by comparing labeling patterns between the dCA1-MEC and ventral CA1 (vCA1)-MEC circuits. In our previous study using channelrhodopsin-2 (ChR2)-assisted circuit mapping, which combines optogenetic stimulation with slice patch-clamp recordings, we demonstrated that the organization of the CA1-MEC circuit differs along the dorsoventral axis: dCA1 neurons predominantly target LVb neurons in the dorsal MEC, whereas vCA1 neurons preferentially target layer Va (LVa) neurons across the entire dorsoventral extent of the MEC (Ohara et al. 2023).

As shown in Fig. 1 and Fig. 2, injection of recombinase-expressing AAVs into the dCA1, together with Coff/Fon-EYFP into the dorsal MEC, resulted in robust labeling of neurons predominantly in LVb of the dorsal MEC (dCA1-recipent MEC neurons). The proportion of labeled neurons in LVb was significantly higher than that in other layers (Fig. 3a-c; LVI, 15.9 ± 4.7%; LVb, 54.9 ± 2.6%; LVa, 18.0 ± 3.5%; LIII, 5.5 ± 0.7%; LII, 5.7 ± 1.9%; LI, 0%; one-way ANOVA, F(5, 18) = 43.5, p < 0.0001, Bonferroni’s multiple comparison test, p < 0.001 for LVb vs. other layers). Among the remaining layers, labeled neurons were preferentially distributed in LVa and layer VI (LVI), although the proportions in these layers were not significantly higher than those in layers II (LII) and LIII.

**Fig. 3.**
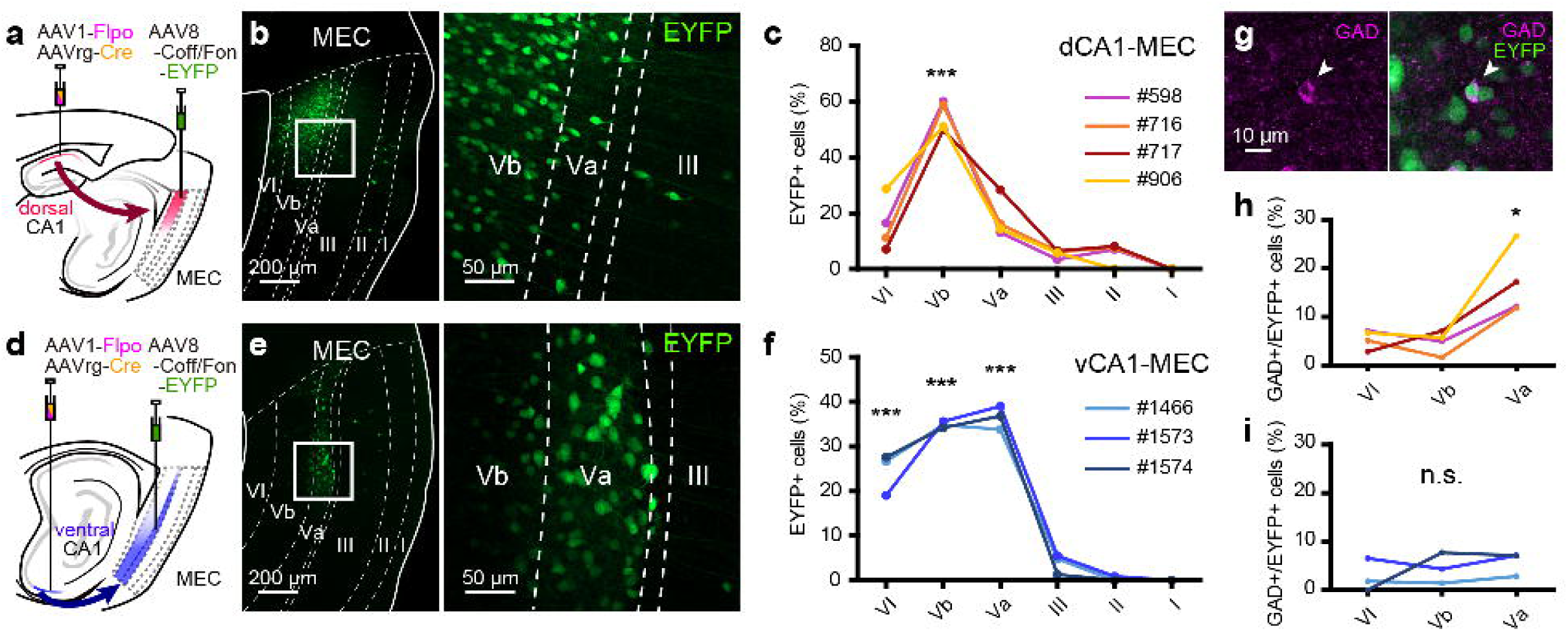
Analysis of the MEC innervation by the dorsal and ventral hippocampal neurons. **a** Schematic diagram of the dCA1-MEC sample. Projections from the dorsal CA1 to MEC were examined by injecting recombinase-expressing AAVs (AAV1-Flpo and AAVrg-Cre) into the dorsal hippocampus and the intersectional AAV (AAV8-Coff/Fon-EYFP) into MEC. **b** Representative fluorescent image of a dCA1-MEC sample showing the laminar distribution of EYFP-labeled neurons in MEC. **c** Proportion of EYFP-positive neurons in each MEC layer among the total labeled neurons in the dCA1-MEC sample (N = 4; one-way ANOVA, F(5, 18) = 43.5, *p* < 0.0001, Bonferroni’s multiple comparison test, \*\*\**p* < 0.001). **d** Schematic diagram of the vCA1-MEC sample. Projections from the ventral CA1 to MEC were examined by injecting recombinase expressing AAVs into the ventral CA1 while injecting the intersectional AAV to MEC. **e** Representative fluorescent image of a vCA1-MEC sample showing the laminar distribution of EYFP-labeled neurons in MEC. **f** Proportion of EYFP-positive neurons in each MEC layer among the total labeled neurons in the vCA1-MEC sample (N = 3; one-way ANOVA, F(5, 12) = 128.3, *p* < 0.0001, Bonferroni’s multiple comparison test, \*\*\**p* < 0.001). **g** Representative fluorescent image of a hippocampus-targeted MEC GABAergic neuron co-labeled with GAD and EYFP (white arrowhead). **h, i** Proportion of GAD-positive neurons among the GFP-positive neurons in layers Va, Vb, and VI in the dCA1-MEC sample (H; one-way ANOVA, F(2, 9) = 11.66, *p* = 0.0086, Bonferroni’s multiple comparison test, * *p* < 0.05) and the vCA1-MEC sample (I; one-way ANOVA, F(2, 6) = 0.86, *p* = 0.48).

In contrast, injection of recombinase-expressing AAVs into the vCA1 resulted in a distinct laminar distribution of labeled neurons in the dorsal MEC. In this condition, labeled neurons were distributed in LVa in addition to LVb (vCA1-recipent MEC neurons) (Fig. 3d-f; LVb, 34.9 ± 0.4%; LVa, 36.5 ± 1.5%), consistent with the known projection patterns of the vCA1 neurons. The proportions of labeled neurons in LVa and LVb were significantly higher than those in layers I-III (one-way ANOVA, F(5, 12) = 128.3, p < 0.0001, one-way ANOVA followed by Bonferroni’s multiple comparison test, p<0.001 for LVb vs layers I–III and LVa vs layers I–III). In addition, a substantial proportion of labeled neurons was detected in LVI (24.4 ± 2.7%), which was significantly higher than that in layers I–III (p < 0.001) but lower than those observed in LVb and LVa (p<0.05 for LVI vs LVb; p < 0.01 for LVI vs LVa). To further examine the difference in labeling patterns between LVa and LVb, we quantified the density of labeled neurons within LVa and LVb in 200-µm columnar bins (Supplementary Fig. 2a; see Methods for detail). The density of labeled neurons was significantly higher in LVa than in LVb (Supplementary Fig. 2b, C; two-tailed paired t test, t2 = 37.58, p = 0.0007).

To further characterize the innervation patterns of dCA1 and vCA1 neurons onto MEC LV/VI neurons, we immunostained the sections for glutamic acid decarboxylase (GAD) to assess the proportion of GABAergic neurons among the CA1-recipent MEC neurons (Fig. 3g). Among dCA1-recipent MEC neurons, the proportion of GAD-positive cells was significantly higher in LVa compared to those of LVb and LVI (Fig. 3h; one-way ANOVA, F(2, 9) = 11.66, p = 0.0086, Bonferroni’s multiple comparison test, p < 0.05 for LVI vs LVa and LVb vs LVa). In contrast, no significant layer-dependent differences in the proportion of GAD-positive neurons were observed among vCA1-recipent MEC neurons (Fig. 3i; one-way ANOVA, F(2, 6) = 0.86, p = 0.48). Together, these results indicate that dCA1 and vCA1 neurons innervate MEC with not only distinct laminar projection patterns but also differential preferences for postsynaptic cell types.

### Whole-brain mapping of the ventral hippocampal output circuits using dsATT with systemic delivery of reporter construct

We next applied dsATT to examine ventral hippocampal output circuits beyond the entorhinal cortex across the whole brain. To achieve whole-brain labeling, we used AAV-PHP.eB, a serotype that efficiently transduces the central nervous system in adult animals (Chan et al. 2017). Specifically, a cocktail of recombinase-expressing AAVs, AAV1-EF1α-Flpo and AAVrg-hSyn-Cre-WPRE, was injected into the ventral subiculum (vSUB), while AAV-PHP.eB-VTKS6-GFP, which expresses GFP in the presence of Flpo and in the absence of Cre (Pouchelon et al. 2022), was administered intravenously (Fig. 4a). We observed transsynaptically labeled neurons across multiple brain regions, including the medial frontal cortex (MFC), nucleus accumbens (NAc), septal region, thalamic nucleus reuniens (RE), anterior hypothalamic nucleus (AHN), basolateral amygdaloid nucleus (BLA), basomedial amygdaloid nucleus (BMA), and the mammillary body (MB) (Fig. 4b-f).

**Fig. 4.**
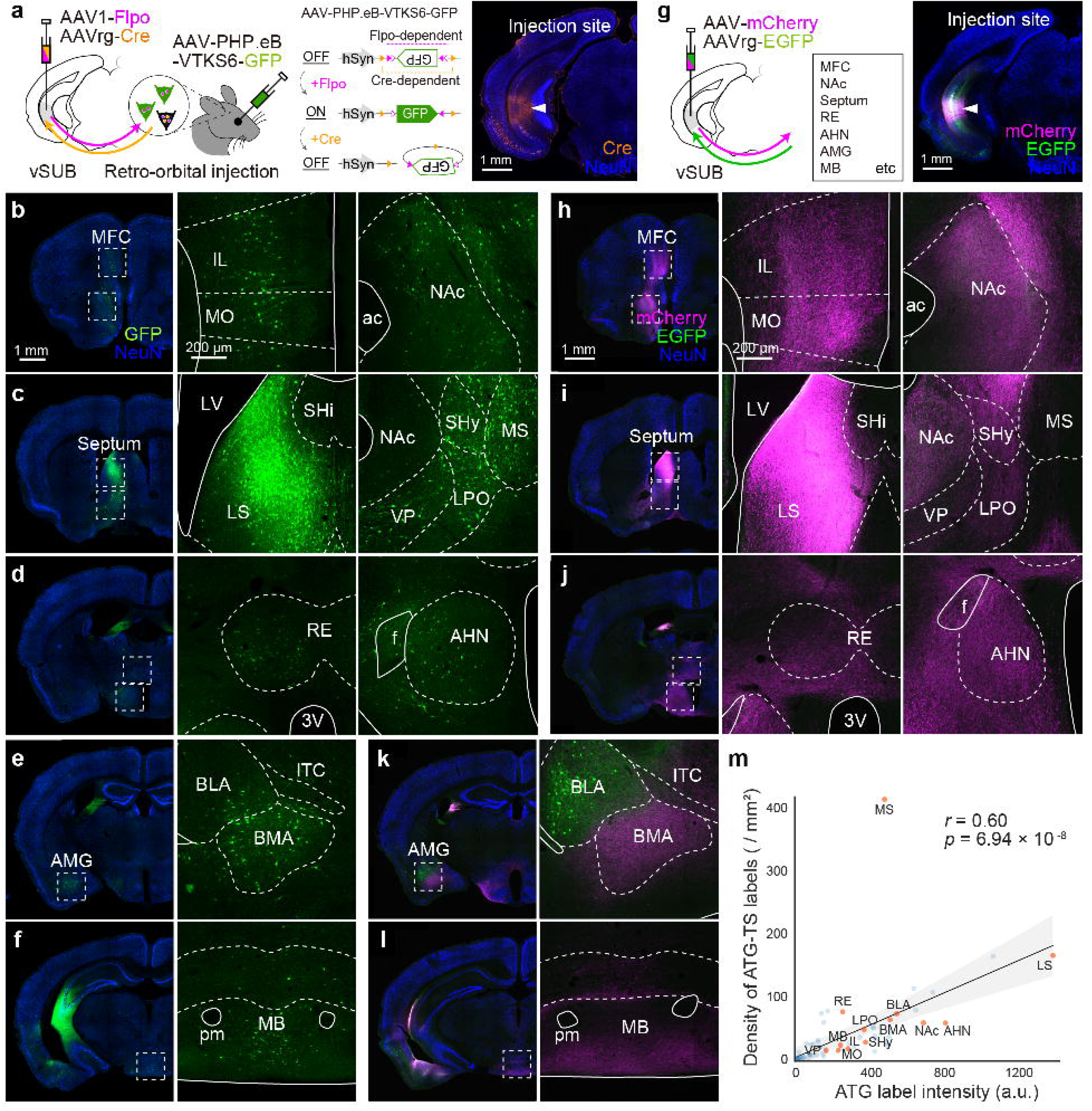
Whole-brain analysis of ventral hippocampal output circuits. **a** Schematic diagram of the anterograde transsynaptic tracing from vSUB with systemic delivery of Coff/Fon construct (left) and representative image of vSUB injection site (right). AAV1-Flpo and AAVrg-Cre were co-injected into vSUB while AAV-PHP.eB-VTKS6-GFP was delivered by retro-orbital injection. **b-f** Representative images showing the distribution of anterograde transsynaptically labeled neurons (green) in MFC, NAc, septal region, RE, AHN, BLA, BMA, and MB. **g** Schematic diagram of the anterograde and retrograde tracing for anatomical reference of vSUB connectivity (left) and the representative image of vSUB injection site (right). Anterograde (AAV-mCherry) and retrograde (AAVrg-EGFP) AAVs were co-injected into vSUB. **h-l** Representative images showing the distribution of anterogradely labeled axons (magenta) and retrogradely labeled neurons (green) in MFC, NAc, septal region, RE, AHN, BLA, BMA, and MB. **m** Positive correlation between the fluorescent intensity of anterograde axonal labels from vSUB (ATG label intensity) and the density of anterograde transsynaptic (ATG-TS) labeling across brain regions (Pearson’s *r* = 0.60, *p* = 6.94 × 10⁻^8^). Each dot represents the values for each brain region, averaged across animals (orange, brain regions shown in b–f and h–l; blue, not shown). The solid line and shaded area indicate the linear regression line and 95% confidence interval, respectively. The medial septum (MS) deviated from the overall trend (standardized residual = 7.44), showed relatively dense ATG-TS labeling despite low projection intensity. ac anterior commissure; AHN, anterior hypothalamic nucleus; BLA, basolateral amygdaloid nucleus; BMA, basomedial amygdaloid nucleus; f, fornix; ITC, intercalated nucleus; IL, infralimbic cortex; LPO, lateral preoptic area; LS, lateral septum; LV, lateral ventricle; MB, mammillary body; MO, medial orbital cortex; MS, medial septum; NAc, nucleus accumbens; pm, principal mammillary tract; RE, thalamic nucleus reuniens; SHi, septohippocampal nucleus; SHy, septohypothalamic nucleus; VP, ventral pallidum.

To confirm whether these labeling patterns reflect the connectivity of the vSUB, we co-injected an anterograde AAV (AAV1-CaMKII-mCherry or AAV9-hSyn-mCherry) together with a retrograde AAV (AAVrg-hSyn-EGFP) into vSUB (Fig. 4g). Anterogradely labeled axons from vSUB neurons were densely distributed in the brain regions where transsynaptically labeled neurons were observed, and the projection patterns closely matched the distribution of the transsynaptically labeled neurons (Fig. 4h-l).

We further examined whether the density of anterograde transsynaptic (ATG-TS) labels in each brain region is correlated with the intensity of axonal projections from vSUB. The density of ATG-TS labels from vSUB was measured using the datasets obtained by anterograde transsynaptic tracing with systemic delivery of VTKS6 reporter by AAV-PHP.eB (n = 3). The intensity of axonal projections from vSUB was obtained from vSUB anterograde tracing datasets (n = 4) and quantified as the mean fluorescence intensity in anterograde labeling channel (ATG label intensity). Comparison of these two values across brain regions revealed a positive correlation between ATG-TS labeling and ATG label intensity (Pearson’s r = 0.60, *p* = 6.94 × 10^−8^; Fig. 4m, Supplementary Table 1), suggesting that abundance of ATG-TS labels likely reflects the projection strength. Notably, linear regression analysis revealed one exception was the labeling in MS (standardized residual = 7.44), where ATG-TS labels were abundant despite sparse axonal projections.

### Analysis of the hippocampal-AMG circuits using dsATT

Among the brain regions targeted by the vSUB, AMG exhibited a distinctive labeling pattern, with anterograde and retrograde labeling distributed topographically across its nuclei (Fig. 4k). To further characterize this organization, we analyzed the distribution of labeling along the anteroposterior axis of AMG (Fig. 5a-d; the same samples as those shown in Fig. 4g-l). To visualize the spatial distribution of anterogradely labeled axons and retrogradely labeled neurons, voxel-based density maps were generated from atlas-aligned sections for each animal and overlaid within a common anatomical reference space (Fig. 5d; see Methods). Consistent with previous reports (Canteras and Swanson 1992; Hintiryan et al. 2021), anterogradely labeled axons originating from vSUB were prominent in the basomedial amygdaloid nucleus, particularly in its posterior part (BMAp). They were also observed in the basolateral amygdaloid nucleus anterior part (BLAa), posterior part (BLAp), and the amygdalohippocampal area (AHi) (Fig. 5b,d). Within these regions, labeling was preferentially distributed toward their posterior portions. In contrast, retrogradely labeled neurons were largely restricted to BLAa and BLAp. These results show that vSUB projections are broadly distributed across multiple amygdala nuclei, including both basolateral and basomedial nuclei, whereas neurons projecting back to the vSUB were predominantly confined to basolateral nuclei.

**Fig. 5.**
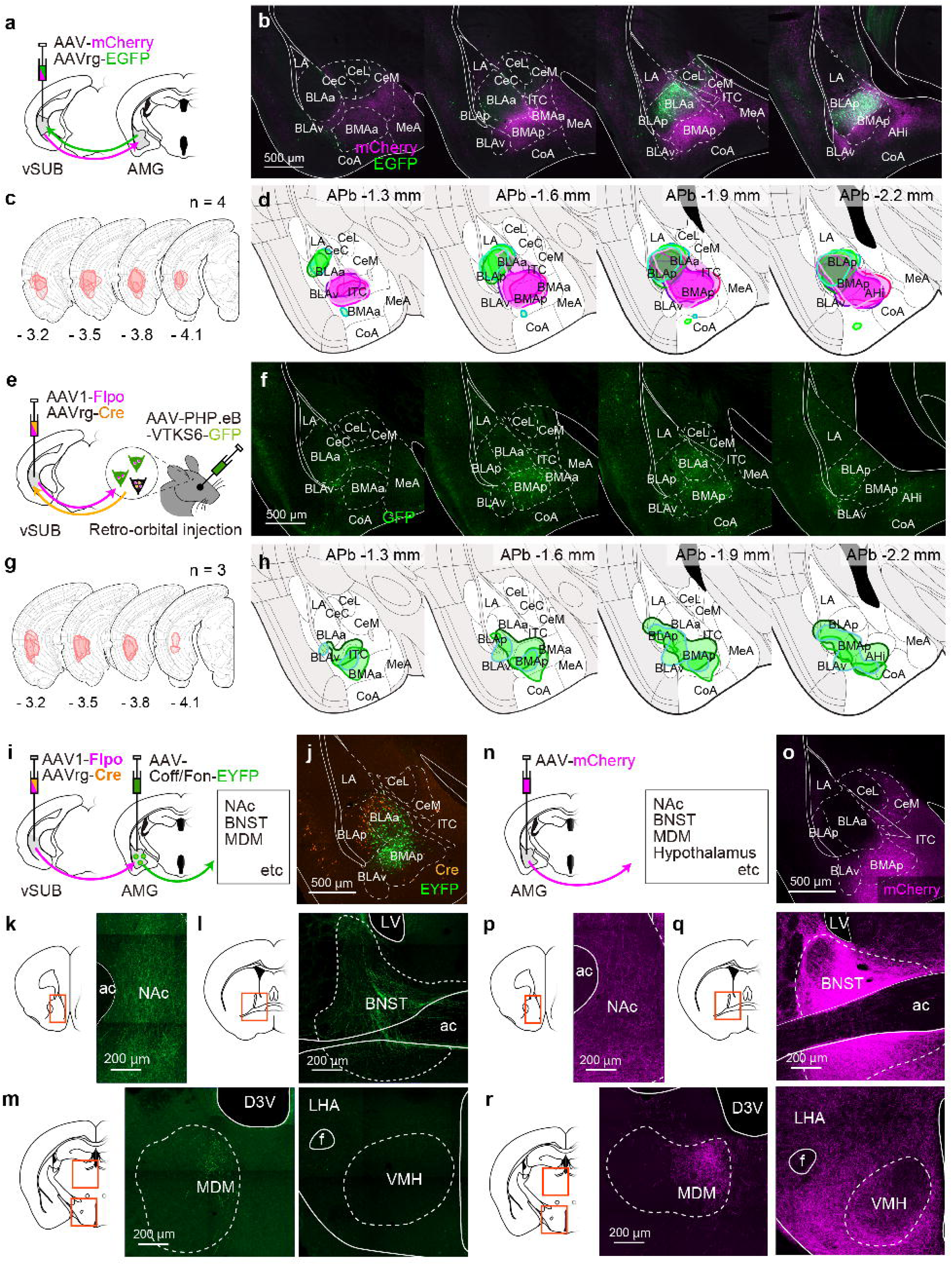
Analysis of the AMG innervation by the ventral subiculum neurons. **a** Schematic diagram of the anterograde and retrograde tracing from vSUB. Anterograde (AAV-mCherry) and retrograde (AAVrg-EGFP) AAVs were co-injected into vSUB. **b, c** Representative amygdala images (the same sample shown in Fig. 4g–l) showing the topographical distribution of anterogradely labeled axons and retrogradely labeled neurons along the antero-posterior axis, and the summary of vSUB injection sites from individual animals superimposed on the coronal sections (n = 4). Numbers in each section give the distance from bregma (APb) in mm. **d** Density map showing the spatial distribution of anterogradely-labeled axons (magenta) and retrogradely labeled neurons (green). Colored areas indicate regions corresponding to the top 1% percentile of smoothed voxel-wise density. Density maps from four animals are overlaid, with different contour colors representing individual animals. **e** Schematic diagram of the anterograde transsynaptic tracing from vSUB with systemic delivery of Coff/Fon construct. AAV1-Flpo and AAVrg-Cre were co-injected into vSUB while AAV-PHP.eB-VTKS6-GFP was delivered by retro-orbital injection. **f, g** Representative amygdala images (the same sample shown in Fig. 4a-f) showing the topographical distribution of anterograde transsynaptically labeled neurons along the antero-posterior axis, and the summary of vSUB injection sites. Numbers in each section give the distance from bregma (APb) in mm. **h** Density map showing the spatial distribution of anterograde transsynaptic labeled neurons. Colored areas indicate regions corresponding to the top 1% percentile of smoothed voxel-wise density. Density maps from three animals are overlaid, with different contour colors representing individual animals. **i** Schematic diagram of the vSUB-AMG sample. Projections from AMG neuronal subpopulations receiving vSUB inputs were examined by injecting recombinase-expressing AAVs (AAV1-Flpo and AAVrg-Cre) into the vSUB and the intersectional AAV (AAV8-Coff/Fon-EYFP) into AMG. **j-m** Representative vSUB-AMG sample. Anterograde transsynaptic labeled AMG neurons (green) were mainly seen in BMAp, and their projections were observed in NAc, BNST, and MDM, but not in the hypothalamus. **n** Schematic diagram of the anterograde tracing for anatomical reference of BMAp projections. AAV9-mCherry was injected into BMAp. **o-r** Representative sample of anterograde tracing from BMAp. The core of injection was mainly in BMAp, and anterogradely labeled axons (magenta) were observed in NAc, BNST, MDM, and also in the hypothalamus. ac, anterior commissure; AHi, amygdalohippocampal area; BLAa/p/v, anterior/posterior/ventral basolateral nucleus; BMAa/p, anterior/posterior basomedial nucleus; BNST, bed nucleus of stria terminalis; CeC/L/M, central nucleus capsular/lateral/medial part; CoA, cortical amygdaloid nucleus; D3V, dorsal third ventricle; f, fornix; ITC, intercalated nucleus; LA, lateral nucleus; LHA, lateral hypothalamic area; LV, lateral ventricle; MeA, medial nucleus; MDM, medial part of mediodorsal thalamic nucleus; NAc, nucleus accumbens; VMH, ventromedial hypothalamic nucleus.

We next examined whether similar spatial patterns were observed using dsATT (Fig. 5e-h; the same samples as those shown in Fig. 4a-f). Transsynaptically labeled neurons were observed in posterior BLAa, BLAp, BMAp, and AHi, largely overlapping with the projection fields of vSUB axons within the amygdala. Notably, BMAp contained a major population of amygdala neurons receiving synaptic input from the vSUB. We next asked whether these vSUB-recipient AMG neurons project to distinct targets. To selectively label these neurons and trace their axons, we performed direct injection of the intersectional viral vector into AMG (Fig. 5i-m). Specifically, a cocktail of AAV1-EF1α-Flpo and AAVrg-hSyn-Cre-WPRE was injected into vSUB, while AAVDJ-hSyn-Coff/Fon EYFP-WPRE was injected into AMG (Fig. 5i). Similar to the retro-orbital injection condition, EYFP-labeled neurons were mainly observed in BMAp (Fig. 5j). EYFP-labeled axons were detected in several extra-amygdaloid regions, including the nucleus accumbens (NAc), the bed nucleus of the stria terminalis (BNST), and the medial part of the mediodorsal thalamic nucleus (MDM) (Fig. 5k-m).

To compare the projections of vSUB-recipient BMAp neurons with those of the bulk BMAp population, anterograde tracing was performed by injecting AAV9-hSyn-mCherry into AMG centered on BMAp (Fig. 5n–r). Anterogradely labeled axons from BMAp neurons were observed in NAc, BNST, and MDM (Fig. 5p-r) Notably, dense labeled axons were also observed in the hypothalamus (Fig. 5r), where axons from transsynaptically labeled BMAp neurons were not detected (Fig. 5m). These observations suggest that the subset of BMAp neurons receiving vSUB input constitutes a distinct neuronal subpopulation with selective projection targets: vSUB neurons preferentially target BMAp neurons projecting to NAc, BNST, and MDM, but not those projecting to the hypothalamus.

## Discussion

Defining postsynaptic neuronal populations that receive direct hippocampal inputs is critical for understanding hippocampal output circuits, yet remains technically challenging. One promising approach is anterograde transsynaptic tracing using AAV1. However, AAV1, while capable of anterograde transsynaptic propagation, is also known to undergo retrograde transport (Tervo et al. 2016; Zingg et al. 2017). Although its retrograde efficiency is lower than that of AAVrg (Zingg et al. 2020), our data indicate that retrograde infection occurs more efficiently than transsynaptic transport to postsynaptic neurons (Fig. 1a-d), thereby limiting its utility for selectively labeling postsynaptic targets. To overcome this limitation, we combined AAV1-mediated anterograde transsynaptic tracing with an intersectional gene expression system. By employing the Coff/Fon construct, we successfully restricted labeling predominantly to the anterograde direction. This approach enables input-defined circuit dissection of hippocampal outputs and allowed us to characterize the detailed organization of hippocampal-MEC and hippocampal-AMG circuits.

### Organization of the hippocampal-MEC circuit

Consistent with our previous study (Ohara et al. 2023), the modified anterograde tracing method revealed laminar differences in MEC neurons targeted by dorsal and vCA1. Specifically, dorsal CA1 preferentially targeted the LVb neurons in MEC, whereas vCA1 innervated a larger proportion of LVa neurons. Interestingly, among MEC neurons targeted by dorsal CA1, the proportion of GABAergic neurons was significantly higher in LVa than in LVb. Our previous study demonstrated that the excitatory/inhibitory neuronal proportion correlates with the excitation/inhibition balance (Alemán-Andrade et al. 2025), suggesting that dorsal CA1 input may induce stronger inhibition in LVa than in LVb of MEC.

Taken together with the preferential targeting of MEC LVb neurons by dorsal CA1, these findings suggest that dorsal CA1 input may excite LVb neurons while indirectly suppressing LVa neurons. This effect is functionally similar to the proposed reciprocal inhibition between LVa and Vb, mediated indirectly via fast spiking interneurons (Rozov et al. 2020). Such circuitry may function as a switch that biases activity toward one of two partially independent layer V circuits in MEC: the “LVa-mediated output circuit”, which relays the hippocampal information to telencephalic regions, and the “LVb-mediated loop circuit” which returns hippocampal signals back to the hippocampus via MEC LIII neurons (Ohara et al. 2018, 2021). This mechanism could help segregate information flow between these partially independent hippocampal-entorhinal circuits, thereby preventing interference between distinct modes of information processing.

### Organization of the hippocampal-AMG circuit

Previous anatomical studies have established dense projections from the ventral hippocampal formation to the amygdala, particularly to medial and basomedial nuclei, as well as reciprocal connectivity between the hippocampus and the basolateral nucleus (Canteras and Swanson 1992; Hintiryan et al. 2021; Pitkänen et al. 2000). Our anterograde and retrograde tracing results are consistent with these reports and further reveal spatial heterogeneity along the antero-posterior axis, with strong reciprocal connectivity in the BLAp and preferential vSUB input to the BMAp.

Our modified anterograde transsynaptic tracing identified a defined population of BMAp neurons receiving direct input from vSUB. Axonal terminals of these input-defined neurons were detected in the extra-amygdaloid regions, including the BNST, MD, and NAc. These regions are widely recognized as key components of extended limbic networks involved in affective regulation, limbic–cortical communication, and motivational processing (Davis et al., 2010; Janak & Tye, 2015; Mitchell, 2015). Notably, projections to the hypothalamus were not detected from this input-defined population, despite known BMAp projections to hypothalamic regions. Indeed, a previous double retrograde tracing study has shown that the amygdala contains distinct subpopulations of neurons projecting to different target regions (Reppucci & Petrovich, 2016). Taken together, our results suggest that hippocampal outputs may preferentially recruit a subpopulation of BMAp neurons involved in integrative and modulatory functions, rather than in direct hypothalamic effector pathways.

### Pros and Cons of dsATT

As demonstrated in the hippocampal-MEC circuit and hippocampal-AMG circuit, dsATT using the Coff/Fon construct enables to examine the detailed circuitry in reciprocally connected brain regions. By further combining this method with systemic delivery of intersectional reporter using AAV-PHP.eB, we succeeded in mapping hippocampal output circuits at the whole brain level. The overall distribution of labeled neurons was largely consistent with the projection patterns identified by anterograde tracing, and the density of transsynaptically labeled neurons positively correlated with the intensity of the axonal projections. This indicates that the quantification of the transsynaptically labeled neurons can be utilized to predict the projection strength.

By introducing genes encoding channelrhodopsin or DREADDs instead of fluorescent proteins, dsATT can be extended to investigate not only organization but also the function of the hippocampal output circuits. Furthermore, because dsATT relies exclusively on viral gene delivery and does not require transgenic lines, it offers broad applicability across species. In principle, this approach can be extended to other mammalian models, including rats and non-human primates such as marmosets and macaques, in which transgenic tools remain limited.

However, as with other approaches using viral vectors, the labeling pattern obtained with this method is constrained by the infection properties of the viruses. AAV2rg, which is widely used as a retrograde AAV, has been reported to exhibit relatively low retrograde infection efficiency to several brain regions such as the MS and thalamus (Han et al., 2023). Indeed, we observed only sparse retrograde labeling in MS (Fig. 4i), even though MS has dense projection to the ventral hippocampus (Nyakas et al., 1987; Ohara et al., 2013). In addition, since the axonal projection from the vSUB to MS was sparse (Fig. 4i), we assume that the transsynaptically labeled neurons observed in MS represent off-target labeling caused by retrograde infection of AAV1-EF1α-Flpo in the absence of AAVrg-hSyn-Cre-WPRE (Fig. 4c). This limitation may be mitigated by using alternative retrograde viruses, such as AAV11, which show efficient retrograde infection of the neurons in MS and the thalamus (Han et al., 2023).

Together, these observations indicate that dsATT provides a practical framework for examining the organization of hippocampal output circuits at the cellular level, while also highlighting the importance of viral tropism in determining labeling specificity. Further improvements in viral vectors may enable more precise identification of input-defined neuronal populations. A recent study has also reported an engineered protein, ATLAS, that propagates across excitatory synapses and enables cell-type specific transsynaptic labeling (Rivera et al., 2025), highlighting the continuing development of molecular tools for circuit-specific tracing. These advances will facilitate systematic dissection of hippocampal output pathways and provide new insights into how hippocampal signals are processed across distributed brain circuits.

## Acknowledgments

We thank Patrick Nicolaas Ang for help with histological preparations and microscopic imaging.

## Author contributions

Conceptualization: S.O.; Investigation: all authors; Supervision: S.O.; Writing, original draft: S.O. and T. K.; Writing, review and editing: all authors; Funding acquisition: S.O., and KI.T.

## Data availability

The data is available from the corresponding author upon request. Source data are provided with this paper.

