## Supplementary information for "Mapping mouse hippocampal output circuits using direction-selective anterograde transsynaptic transduction"

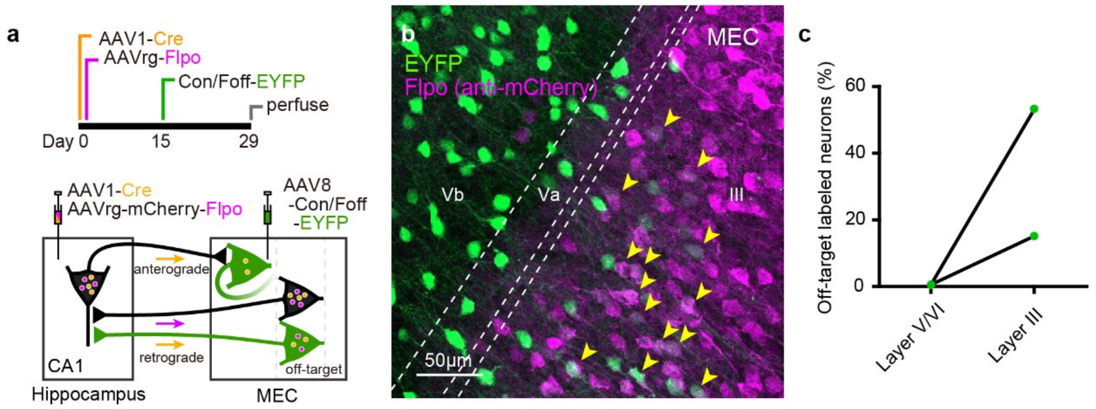


**Supplementary Figure 1. Off-target labeling with the Con/Coff construct after delayed injection.**

**a** Schematic diagram of dsATT illustrating the timing of the viral injections. AAV8-Con/Foff-EYFP was injected into MEC 15 days after the injection of an AAV cocktail (AAV1-Cre and AAVrg-mCherry-Flpo) into the dorsal CA1. **b** Representative fluorescent image of MEC. Yellow arrowheads indicate off-target labeled neurons co-expressing EYFP and mCherry-Flpo. **c** Proportion of off-target labeled neurons in layer III and in deep layers (V/VI) of MEC (N = 2).

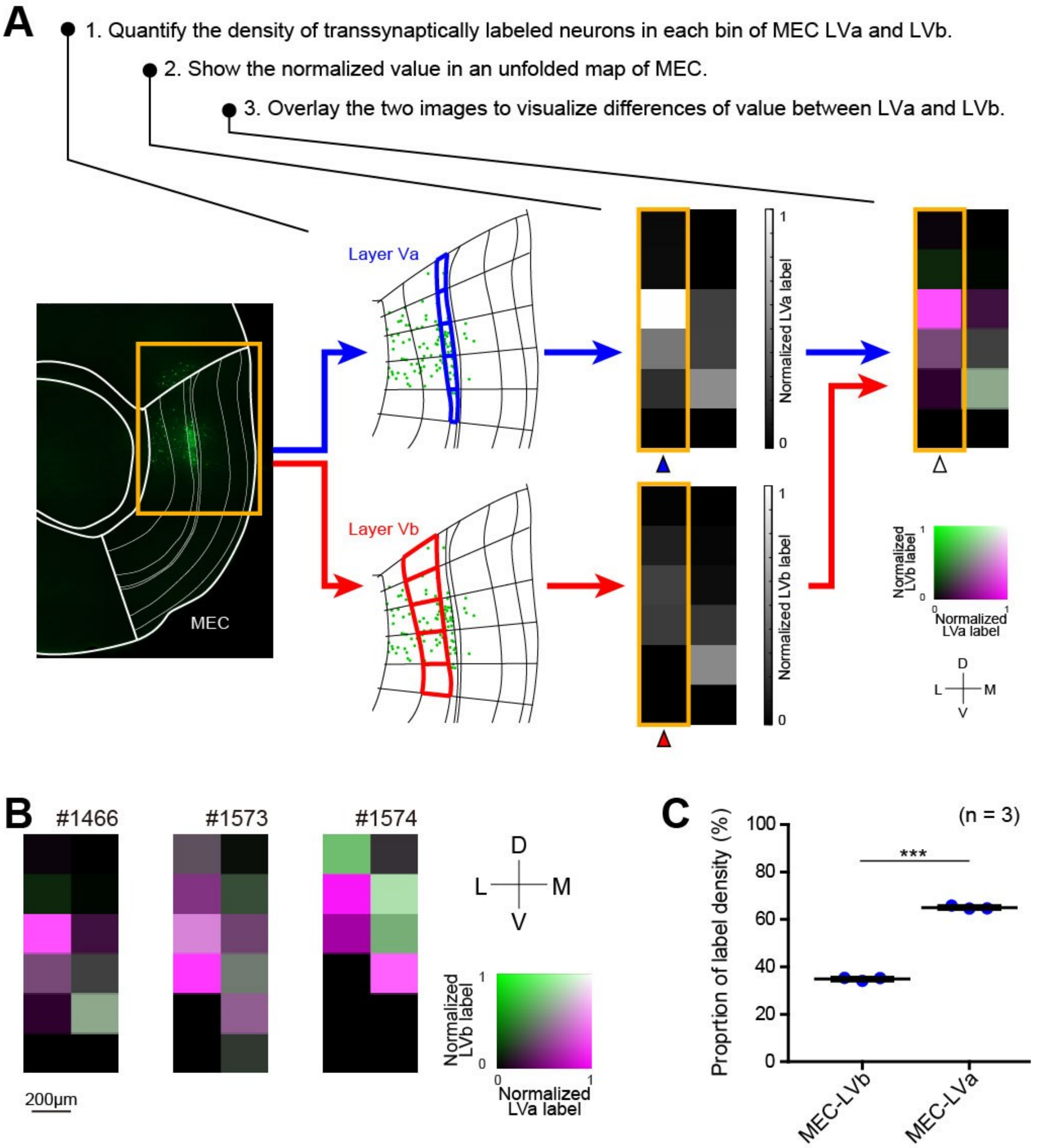


**Supplementary Figure 2. Density of transsynaptically labeled neurons in MEC layers Va and Vb.**

**a** Schematic diagram illustrating the quantitative analysis of the density of transsynaptically labeled neurons in EC layers Va and Vb. Layers Va and Vb were divided into 200-µm columnar bins and the density of labeled neurons of each bin was quantified (step 1). The intensity of each bin was then normalized for every sample and the normalized intensities were mapped on an unfolded map of MEC in grayscale (step 2). Lastly, a composite image was created by overlaying these two grayscale maps of LVa and LVb (step 3). In this composite image, differences of labeling patterns between LVa and LVb are shown in different color bands. Magenta indicates bins with a dense labeling of axons in LVa and green indicates bins with densely labeled axons in LVb. Bins with a dense labeling of axons in both LVa and LVb are shown in white and bins with no labeled axons are shown in black. **b** Two-dimensional density maps showing the patterns of transsynaptically labeled neurons in MEC for the three samples (the same samples as those shown in Fig. 3d–f). **c** Proportion of label density in MEC-LVa and MEC-LVb for samples (N = 3; two-tailed paired t test, t2 = 37.58, *** $p = 0.0007$).

**Table S1. Original data of the linear regression analysis of whole-brain labeling patterns shown in Figure 4M.**

| Abbreviation | Name | ATG label intensity (a.u.) (N = 4) | | ATG-TS label density (/mm$^2$) (N = 3) | |
|---|---|---|---|---|---|
| | | Mean | SD | Mean | SD |
| M1 | Primary motor cortex | 7.53 | 1.32 | 3.77 | 0.24 |
| M2 | Secondary motor cortex | 12.95 | 3.32 | 3.36 | 1.30 |
| S1 | Primary somatosensory cortex | 11.50 | 1.63 | 3.64 | 0.83 |
| S2 | Secondary somatosensory cortex | 9.62 | 3.52 | 3.01 | 0.70 |
| A24a (IL) | Cingulate cortex, area 24a (Infralimbic cortex) | 289.59 | 85.29 | 19.51 | 18.27 |
| AuD | Secondary auditory cortex, dorsal area | 22.14 | 10.31 | 14.03 | 19.77 |
| AuV | Secondary auditory cortex, ventral area | 33.15 | 21.24 | 3.82 | 2.57 |
| V2L | Secondary visual cortex, lateral area | 61.72 | 57.14 | 27.93 | 20.34 |
| V2M | Secondary Visual Cortex, medial area | 18.06 | 13.79 | 2.95 | 1.20 |
| V1 | Primary visual cortex | 20.92 | 12.86 | 3.17 | 0.34 |
| A24b (Cg1) | Cingulate cortex, area 24b | 49.30 | 25.40 | 5.37 | 2.42 |
| A32 (PrL) | Cingulate cortex, area 32 (Prelimbic cortex) | 251.51 | 122.53 | 19.68 | 22.63 |
| LO | Lateral orbital cortex | 21.28 | 16.49 | 3.51 | 1.32 |
| MO | Medial orbital cortex | 237.21 | 105.29 | 16.78 | 20.46 |
| VO | Ventral orbital cortex | 82.21 | 45.47 | 10.57 | 9.42 |
| AID | Agranular insular cortex, dorsal part | 10.53 | 8.14 | 2.89 | 1.05 |
| A30 (RSD) | Cingulate cortex, area 30 (Retrosplenial dysgranular cortex) | 16.44 | 8.66 | 3.61 | 0.78 |
| A29 (RSG) | Cingulate cortex, area 29 (Retrosplenial granular cortex) | 44.48 | 46.52 | 5.20 | 0.70 |
| TeA | Temporal association cortex | 75.01 | 54.36 | 5.40 | 5.18 |
| PRh | Perihinal cortex | 356.65 | 269.59 | 37.16 | 50.56 |
| Ect | Ectorhinal cortex | 168.08 | 141.91 | 18.70 | 25.95 |
| MOB | Main olfactory bulb | 22.13 | 6.50 | 6.95 | 2.36 |
| AO | Anterior olfactory area | 429.71 | 141.30 | 52.61 | 64.81 |
| A25 (DP) | Cingulate cortex, area 25 (Dorsal peduncular cortex) | 513.44 | 185.71 | 38.15 | 38.27 |
| PIR | Piriform cortex | 85.38 | 48.07 | 10.40 | 6.48 |
| PCO | Posterior cortical amygdaloid area | 139.18 | 36.19 | 22.60 | 11.22 |
| RAPir | Rostral amygdalopiriform area | 101.05 | 37.03 | 26.75 | 10.76 |
| Apir | Amygdalopiriform transition area | 553.11 | 164.12 | 89.60 | 31.15 |
| DLEnt | Dorsolateral entorhinal cortex | 547.48 | 240.91 | 74.54 | 60.53 |
| DIEnt | Dorsal intermediate entorhinal cortex | 753.82 | 161.42 | 109.33 | 58.48 |
| VIEnt | Ventral intermediate entorhinal cortex | 660.89 | 102.04 | 80.86 | 65.42 |
| MEnt | Medial entorhinal cortex | 650.02 | 457.97 | 115.38 | 79.94 |
| CEnt | Caudomedial entothinal cortex | 158.41 | 121.72 | 61.33 | 45.27 |
| PaS | Parasubiculum | 145.73 | 90.87 | 75.56 | 66.96 |
| En | Endopiriform nucleus | 152.83 | 53.01 | 11.03 | 11.84 |
| BLA | Basolateral amygdalar nucleus | 557.88 | 84.25 | 75.24 | 36.12 |
| BMA | Basomedial amygdalar nucleus | 520.95 | 91.43 | 65.53 | 25.88 |
| AHi | Amygdalohippocampal area | 469.21 | 167.70 | 43.37 | 25.59 |
| STRd | Striatum dorsal region | 25.13 | 8.60 | 2.91 | 1.85 |
| NAc | Nucleus accumbens | 702.97 | 69.79 | 60.72 | 28.23 |
| IPAC | Interstitial nucleus of the posterior limb of the anterior commissure | 98.34 | 33.73 | 8.02 | 9.27 |
| Tu | Olfactory tubercle | 119.50 | 34.50 | 9.14 | 3.15 |
| LS | Lateral septal nucleus | 1410.53 | 284.63 | 167.76 | 133.19 |
| SFi | Septofimbrial nucleus | 1083.95 | 270.73 | 166.48 | 106.06 |
| SHy | Septohypothalamic nucleus | 386.54 | 247.73 | 29.99 | 44.69 |
| AA | Anterior amygdaloid area | 78.18 | 41.37 | 10.40 | 4.52 |
| Me | Medial amygdaloid nucleus | 157.96 | 55.25 | 7.35 | 0.54 |
| VP | Ventral pallidum | 171.04 | 78.49 | 16.40 | 13.55 |
| SI | Substantia innominata | 126.43 | 21.74 | 33.12 | 13.95 |
| MS | Medial septal nucleus | 490.19 | 86.99 | 416.24 | 107.46 |
| NDB | Diagonal band nucleus | 127.82 | 25.78 | 30.34 | 18.85 |
| TS | Triangular septal nucleus | 179.80 | 156.88 | 79.77 | 66.86 |
| STa | Bed nuclei of the stria terminalis, anterior division | 505.22 | 147.67 | 24.74 | 13.47 |
| STm | Bed nuclei of the stria terminalis, medial division | 503.96 | 126.39 | 28.76 | 37.14 |
| STl | Bed nuclei of the stria terminialis, lateral division | 435.76 | 99.28 | 40.12 | 28.69 |
| STIA | Bed nucleus of the stria terminalis, intraamygdaloid | 435.00 | 112.65 | 34.84 | 21.12 |
| AV | Anteroventral thalamic nucleus | 87.64 | 78.00 | 26.15 | 33.41 |
| PT | Parataenial thalamic nucleus | 427.15 | 71.36 | 53.61 | 36.73 |
| RE | Reuniens area | 261.59 | 111.89 | 78.14 | 43.72 |
| Rt | Reticular nucleus, prethalamus | 28.60 | 9.53 | 5.80 | 2.20 |
| MPA | Medial preoptic area | 436.16 | 45.86 | 15.01 | 8.59 |
| AHN | Anterior hypothalamic nucleus | 823.26 | 113.83 | 60.43 | 10.72 |
| MB | Mammillary body | 249.70 | 221.63 | 25.12 | 28.02 |
| VMH | Ventromedial hypothalamic nucleus | 323.37 | 68.98 | 22.95 | 6.05 |
| PHA | Posterior hypothalamic area | 248.05 | 87.44 | 40.56 | 4.84 |
| LH | Lateral hypothalamic area | 295.60 | 61.86 | 27.01 | 11.41 |
| LPO | Lateral preoptic area | 378.86 | 84.29 | 50.03 | 38.73 |
| ZI | Zona incerta | 48.77 | 20.94 | 6.17 | 1.80 |

Brain regions were defined according to the Kim laboratory unified mouse brain atlas. Only regions meeting the predefined criteria were included in the analysis (see Methods). ATG intensity represents the mean fluorescence intensity of anterogradely labeled axons in each region after background subtraction. ATG-TS density represents the areal density of neurons labeled by anterograde transsynaptic tracing. Mean values are visualized using a heat map color scale, with higher values indicated by darker red.
